# A test of the Baldwin Effect: Differences in both constitutive expression and inducible responses to parasites underlie variation in host response to a parasite

**DOI:** 10.1101/2020.07.29.216531

**Authors:** Lauren Fuess, Jesse N. Weber, Stijn den Haan, Natalie C. Steinel, Kum Chuan Shim, Daniel I. Bolnick

## Abstract

Despite the significant effect of host-parasite interactions on both ecological systems and organism health, there is still limited understanding of the mechanisms driving evolution of host resistance to parasites. One model of rapid evolution, the Baldwin Effect, describes the role of plasticity in adaptation to novel conditions, and subsequent canalization of associated traits. While mostly applied in the context of environmental conditions, this theory may be relevant to the evolution of host resistance to novel parasites. Here we test the applicability of the Baldwin Effect to the evolution of resistance in a natural system using threespine stickleback fish (*Gasterosteus aculeatus*) and their cestode parasite *Schistochephalus solidus*. We leverage a large transcriptomic data set to describe the response to *S. solidus* infection by three different genetic crosses of stickleback, from a resistant and a tolerant population. Hosts mount a multigenic response to the parasite that is similar among host genotypes. In addition, we document extensive constitutive variation in gene expression among host genotypes. However, although many genes are both infection-induced and differentially expressed between genotypes, this overlap is not more extensive than expected by chance. We also see little evidence of canalization of infection-induced gene expression in the derived resistant population. These patterns do not support the Baldwin Effect, though they illustrate the importance of variation in both constitutive expression and induced responses to parasites. Finally, our results improve understanding of the cellular mechanisms underlying a putative resistance phenotype (fibrosis). Combined, our results highlight the importance of both constitutive and inducible variation in the evolution of resistance to parasites, and identify new target genes contributing to fibrosis. These findings advance understanding of host-parasite interactions and co-evolutionary relationships in natural systems.

## INTRODUCTION

Host-parasite interactions are key drivers of both ecological dynamics (Lefevre et al., 2009) and evolution (Van Valen, 1973;Duffy and Forde, 2009). Furthermore, parasitic infections pose a significant health challenge in a range of vertebrate systems (Thompson et al., 2009;Chomicz et al., 2016). Despite the impact of parasitic infections on both ecosystem function and organismal health, understanding of genetic mechanisms driving rapid evolution of resistance to parasites is limited (Ebert, 2018). Many hypotheses and models of the mechanisms driving rapid evolution have been postulated (Thompson, 1998;Salamin et al., 2010;Kopp and Matuszewski, 2014). One such model is the Baldwin Effect, which describes the role of phenotypic plasticity in enabling the colonization of new habitats and subsequent adaptation to those environments (Baldwin, 1896;Crispo, 2007).

The Baldwin Effect postulates that beneficial traits allowing for adaptation to a novel condition first arise as plastic responses to a variable environment. When the population encounters a new environmental condition due to environmental change or colonization of a new location, plasticity facilitates improved performance by allowing the population to express a new phenotype, perhaps preventing extinction. Subsequently, if plasticity is costly and the new environment is consistent, then selection may favor individuals who constitutively express that new phenotype (e.g., canalization; (Baldwin, 1896). This may entail a loss of plasticity in the derived population, or a shift in the baseline trait while plasticity persists. The Baldwin Effect has been supported by a diversity of empirical studies (Chapman et al., 2000;Adams and Huntingford, 2004;Corl et al., 2018). While most often applied to evolutionary adaptation to new abiotic environments (Yeh and Price, 2004;Badyaev, 2009), the Baldwin Effect itself is likely applicable to adaptation to any new environmental factor, including potential biotic conditions such as a novel parasite. Still, to our knowledge, the Baldwin Effect has yet to be applied to questions of rapid evolution in the context of immunology or host-parasite dynamics.

While the Baldwin Effect itself has yet to be applied to the evolution of host resistance to parasites, there is evidence to suggest that both plastic (i.e. induced) and canalized (i.e. constitutive) differences are important factors driving variation in host resistance. Plastic or induced responses are known to contribute to variation in parasite resistance in both vertebrate and invertebrate systems (Wakelin and Donachie, 1983;Reeson et al., 1998). In contrast, other studies have documented the importance of constitutive variation in parasite resistance (Evison et al., 2016;Kamiya et al., 2016). In truth, enhanced parasite resistance is likely the result of a combination of induced and constitutive traits (Hamilton et al., 2008). Still, while the importance of both sources of variation has been highlighted, the mechanisms producing these differences, and their evolutionary trajectories, remain unknown. In particular: are infection-induced traits also the same phenotypes that later evolve constitutive differences between populations, as the Baldwin Effect would predict? Also, are these constitutive differences between populations a result of canalization (loss of plasticity) in the derived population, or do they rather reflect a shift in baseline traits while plasticity remains?

Here we test the applicability of the Baldwin Effect to the evolution of host resistance, using a powerful natural model system, the threespine stickleback, *Gasterosteus aculeatus*. Sticklebacks’ evolutionary history has yielded extensive natural variation in host-parasite interactions that can be used to address questions pertaining to the evolution of host resistance. Ancestral marine stickleback repeatedly colonized freshwater environments during the Pleistocene deglaciation, resulting in replicated, independent evolution (McKinnon and Rundle, 2002). Colonization of freshwater lakes exposed stickleback to a helminth cestode parasite, *Schistocephalus solidus,* which is absent in marine environments (Simmonds and Barber, 2016) and rare in anadromous stickleback that spend a small portion of their life in fresh or brackish water. Consequently, numerous freshwater populations of stickleback have been independently evolving in response to this freshwater-associated parasite for thousands of generations, creating a system of repeated evolution of increased resistance to *S. solidus* relative to still-extant marine populations (Weber et al., 2017a). However, the resulting host resistance varies among lake populations. In many lakes stickleback are less likely to be infected (compared to marine fish), but when infected they allow rapid parasite growth to >30% of host body mass. Yet in some populations stickleback evolved an additional capacity to severely reduce cestode growth and can eliminate the parasite (Weber et al., 2017b). Preliminary findings indicate that the growth suppression in these lakes is caused, in part, by the formation of fibrotic scar tissue that traps the parasite (Weber et al., *in prep*), and can sometimes kill the parasite. This among-lake variation in parasite resistance and fibrosis presents a powerful system for investigating mechanisms of host resistance and the processes controlling their evolution, with a clear delineation between which traits are ancestral (marine, or growth-permitting) and derived (fibrotic and growth-suppressing).

Here we report findings from transcriptomic analysis of *G. aculeatus* experimentally exposed to *S. solidus*, and use these results as a test of the applicability of the Baldwin Effect to evolution of host resistance in a natural host-parasite system. To that end we test for three kinds of gene expression differences. First, we determine the plastic host response to the parasites’ presence/absence, common to all host genotypes. Second, we measure constitutive differences in gene expression among genetically divergent crosses between two lake populations (one resistant with high fibrosis, the other susceptible with low fibrosis). If the Baldwin Effect is true, we expect high overlap between the genes in these two sets of results. The third test is for genes that are more plastic in response to infection in one genotype than others (an infection by genotype interaction). If canalization contributes to the Baldwin Effect, we additionally expect the derived (resistant) population to exhibit reduced gene expression plasticity compared to the susceptible population, but shifted in the direction of infection-induced expression.

In addition to testing for the Baldwin Effect, we also document the immunogenetic mechanisms associated with a putative parasite resistance phenotype, fibrosis (Weber et al., *in prep*). Finally, we test for signatures of counter-evolution by the parasite by assessing signatures of parasite interference with host responses. Combining these avenues of research we provide improved understanding of host response to parasites and the evolution of host resistance that can increase understanding of host-parasite evolutionary dynamics.

## METHODS

### Experimental Design

We used minnow traps to capture reproductively mature stickleback from Roberts Lake and Gosling Lake, on Vancouver Island in British Columbia. These populations represent two ends of the natural spectrum of parasite prevalence: high parasite load in Gosling, low in Roberts (Weber et al., 2017b). Furthermore, data suggests that Roberts Lake fish have evolved a fibrosis-based immune response to suppress parasite growth, which is not present in Gosling Lake fish (Weber et al. *in prep*). Wild-caught gravid females were stripped of eggs, which we fertilized using sperm obtained from macerated testes of males from the same lake (within-population crosses, denoted ROB or GOS) or the other lake (F1 hybrids, RG or GR depending on cross direction). Fish were collected with permission from the Ministry of Forests, Lands, and Natural Resource Operations of British Columbia (Scientific Fish Collection permit NA12-77018 and NA12-84188). The resulting eggs were shipped back to Austin, Texas, hatched, and reared to maturity. A subset of these first-generation lab-raised adults were experimentally infected with *Schistocephalus solidus* cestodes, or sham-exposed as a control. The resulting infection rates, cestode growth rates, and host immune traits are reported in Weber et al 2017, and host immune gene expression is described in Lohman et al (2017). The remaining lab-reared adults were artificially crossed to generate F2 hybrids, including both intercrosses (F1xF1 hybrids), and reciprocal backcrosses (ROBxF1 or GOSxF1).

We experimentally exposed one-year-old F2 hybrids to *S. solidus* cestodes, following standard procedures (Weber et al., 2017a;Weber et al., 2017b). Briefly, we obtained mature cestodes from wild-caught stickleback from Gosling Lake or Echo Lake (Roberts Lake fish do not carry mature cestodes). We obtained the cestodes by dissecting freshly euthanized fish, then paired the cestodes by mass to mate them in nylon biopsy bags in artificial media (mimicking bird intestines where the cestodes typically mate; (Wedekind et al., 1998). We collected the resulting eggs, and stored these at 4C for up to one year. We hatched the eggs and fed them to *Macrocyclops albidus* copepods. The copepods were screened for successful infections after 14 days, then fed to individually-isolated stickleback, as described in (Weber et al., 2017a;Weber et al., 2017b). We filtered the water after the exposure trial to ensure the copepods had been consumed. All F2 hybrid stickleback used in this trial were exposed to *S. solidus* (no sham exposures), to maximize infection rate for QTL mapping that will be described elsewhere (Weber et al. *in prep*). However, only a subset of fish were successfully infected, providing a contrast between infected versus uninfected fish. Prior transcriptomic and flow cytometry data suggests that fish with failed infections are phenotypically similar to sham exposed fish. The experimentally infected fish were maintained for 42 days post-exposure, then euthanized with MS-222 and dissected to obtain **(1)** one head kidney (pronephros) for flow cytometry; **(2)** one head kidney for gene expression analysis, preserved in RNAlater at −80C; **(3)** fish mass and length and sex; **(4)** the mass and number of successfully established cestodes, and **(5)** the presence or absence of fibrosis. We exposed a total of 711 stickleback to *S. solidus*. All fish handling was approved by the University of Texas IACUC (AUP-2010-00024).

### Flow Cytometry

Flow cytometry data on head kidney cell population ratios (granulocytes versus lymphocytes) and activity (baseline ROS and oxidative burst) was generated following methods described by Weber et al. (Weber et al., 2017a;Weber et al., 2017b). Data was analyzed using FlowJo software (Treestar). Populations of granulocytes and lymphocytes were separated by linear forward scatter (FSC) and side scatter (SSC), providing counts of the relative abundance of each cell type. ROS production by granulocytes was measured following protocols for PMA stimulation described in Weber et al. (Weber et al., 2017a;Weber et al., 2017b).

### RNA extraction and Transcriptome Sequencing

We extracted RNA from one head kidney using the Ambion MagMAX-96 Total RNA Isolation Kit, following a modified version of the manufacturer’s protocol (see supplementary materials). Each head kidney (hereafter ‘sample’) was separately placed in lysis/binding solution and homogenized using a motorized pestle. After initial purification using magnetic beads provided by the kit, DNA was removed by adding TURBO DNAse and a second purification with Serapure magnetic beads, leaving only RNA. The RNA yield of each sample was quantified using a Tecan NanoQuant Plate.

RNAseq libraries were constructed using TagSeq methodologies detailed in Lohman et al. (Lohman et al., 2016) with modifications. After fragmentation of the RNA in a magnesium buffer (NEB Next RNA fragmentation buffer), the RNA fragments were purified using Agencourt RNAClean XP beads. A poly-dT primer (3ILL-30TV) was annealed to the poly-A tail of mRNAs, after which the first cDNA strand was synthesized, which was amplified in a second PCR reaction. The PCR products were purified with Serapure magnetic beads, quantified (Quant-IT PicoGreen) and normalized (1 ng / μL), after which all libraries were PCR-barcoded using Illumina i5 and i7 indexes. Fragment size selection occurred via automated gel extraction and final quantification was performed using Qubit 2.0. The libraries were sequenced using a HiSeq 2500 at the Genomics Sequencing and Analysis Facility of the University of Texas at Austin.

### Bioinformatic Analyses

We processed TagSeq reads (PCR duplicates removed, adaptors trimmed, low quality reads removed) using the iRNAseq pipeline.(Dixon et al., 2015) Reads were aligned to version 95 of the stickleback transcriptome from Ensembl with Bowtie 2 (Langmead and Salzberg, 2012). Samples with less than 500,000 mapped reads were removed from subsequent analyses, resulting in a final *n* = 393. Finally, we annotated transcripts with a blastx comparison to the UniProtKB database (http://www.uniprot.org/help/uniprotkb) with the parameters: max target seqs = 10; evalue = 1e^−5^. Results were filtered to obtain the match with the highest evalue and bit score for each transcript.

### Analysis with DESeq2

To test for differential expression, we used the R package DESeq2. Transcripts were filtered to remove those that were not expressed in more than 195 samples (roughly half of the sample set). The remaining 15,354 sequences were tested for differential expression using the following model:

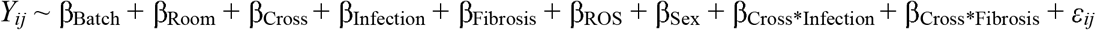

where *Y*_*ij*_ is the count of transcript *i* in individual *j*, β_Room_ is a fixed effect with two levels corresponding to the room in which fish were reared, β_Cross_ is a fixed effect with three levels: F2, GBC, and RBC, *β*_Infection_ is a fixed effect with two levels: infected or uninfected, *β*_Fibrosis_ is a fixed effect with two levels: fibrotic or nonfibrotic, *β*_ROS_ is a continuous factor corresponding to measure reactive oxygen production per sample, and *β*_Sex_ is a fixed effect with three levels: male, female, or unknown (for the few samples where sex could not be identified with confidence). β_Batch_ is a random effect corresponding to the lane of sequence sampling. All *p*-values were multiple test corrected using a 10% FDR (Benjamini–Hochberg; (Benjamini and Hochberg, 1995).

### Expression Pathway and Upstream Regulator Analyses

Differentially activated biological pathways and upstream regulators were assessed for model factors of interest using the Ingenuity Pathway Analysis software (IPA; QIAGEN Inc., https://www.qiagenbioinformatics.com/products/ingenuity-pathway-analysis). For each factor of interest, logfold change, unadjusted p-value, and annotation (spID) per transcript was input into the software. Transcripts were then filtered to retain only those with unadjusted *p*-values < 0.05. Transcripts with duplicated IDs were averaged for analysis. Pathway analyses were used to identify pathways significantly affected by each factor, and generate corresponding *p*-values and activation scores (*z*-scores). Factors tested using IPA were infection, fibrosis, and cross.

### Tests of the Baldwin Effect

The Baldwin Effect makes two predictions that we test here. First, the genes with infection-induced expression changes (plasticity) are also the ones that contribute to constitutive between-population divergence in resistance. We tested this prediction in two ways. We used a χ^2^-test to evaluate whether there is an excess of genes that are significant for both main effects of infection, and main effects of genotype. Then, for the genes that were significant for both effects, we tested for a correlation between their infection and genotype effect sizes: are genes up-regulated by infection also more highly expressed in fish with greater resistant Roberts Lake ancestry? The second prediction is that we should see genotype by infection interactions, in which one genotype is more responsive to infection than the other. Specifically, we expect that a disproportionate fraction of interaction effects entail higher plasticity in the susceptible Gosling Lake genotypes than in fish with greater Roberts lake ancestry. Moreover, the derived resistant genotypes should more closely resemble the infection-induced expression levels. To test we split samples by cross type and ran simplified models (*Y*_*ij*_ ~ β_Batch_ + β_Infection_ + β_ROS_ + β_Sex_ + β_Cross*Infection_ + *ε*_*ij*_) to test for response to infection in RBC and GBC fish separately. We then compared the number of genes which were significantly effected by interaction effects in the full model that also responded significantly to infection in RBC and GBC fish alone. Due to the small number of interaction responsive genes, we did not statistically test whether derived resistance genotypes resemble infection-induced expression levels.

## RESULTS AND DISCUSSION

### Signatures of host response to a common cestode parasite

Host response to infection by a common cestode parasite, *S. solidus*, involved multiple genes and pathways representative of a diversity of immune components. This is a common pattern across systems: host response to parasite often involves multiple arms of the immune system (Anthony et al., 2007;Medzhitov, 2007;Langhorne et al., 2008). Comparing 158 infected versus 232 uninfected fish (all three cross types), we found 2,369 differentially expressed transcripts (*p_adj_* < 0.10, 10% FDR; **Supplementary File 1**), 2,223 of which were annotated. Of these, 341 transcripts were annotated to encode for proteins involved in a range of types of immunity, including antiviral responses (Zinc finger CCCH-type antiviral protein 1; Hayakawa et al., 2010), T-cell functioning (C-type lectin domain family 4 member E; Lu et al., 2018), and Toll-like receptor signaling (Toll-like receptor 8; Cervantes et al., 2012). Biological pathway analyses also indicated broad effects of infection on hosts. A total of 169 pathways were significantly activated as a result of infection (*p_adj_* < 0.10, **Supplementary File 2**). Thirty-one of these pathways are linked to immunity (**Figure 1**). All of these significant immune pathways were suppressed (lower relative transcript abundance) in infected fish relative to uninfected fish. This included a number of pathways involved in inflammation and chemotaxis such as IL-8 signaling (Harada et al., 1994), PPARα/RXRα activation pathway (Youssef and Badr, 2004), and CXCR4 signaling (Stein et al., 2009). Finally, a number of pathways related to phagocytosis were suppressed including Fcy receptor-mediated phagocytosis in macrophages and monocytes, phagosome formation, and phagosome maturation. Overall, these results suggest a broad tendency for key components of immune response to be suppressed in infected fish, including inflammation and immune cell chemotaxis, as well as immune effector processes such as phagocytosis.

**Figure 1:**
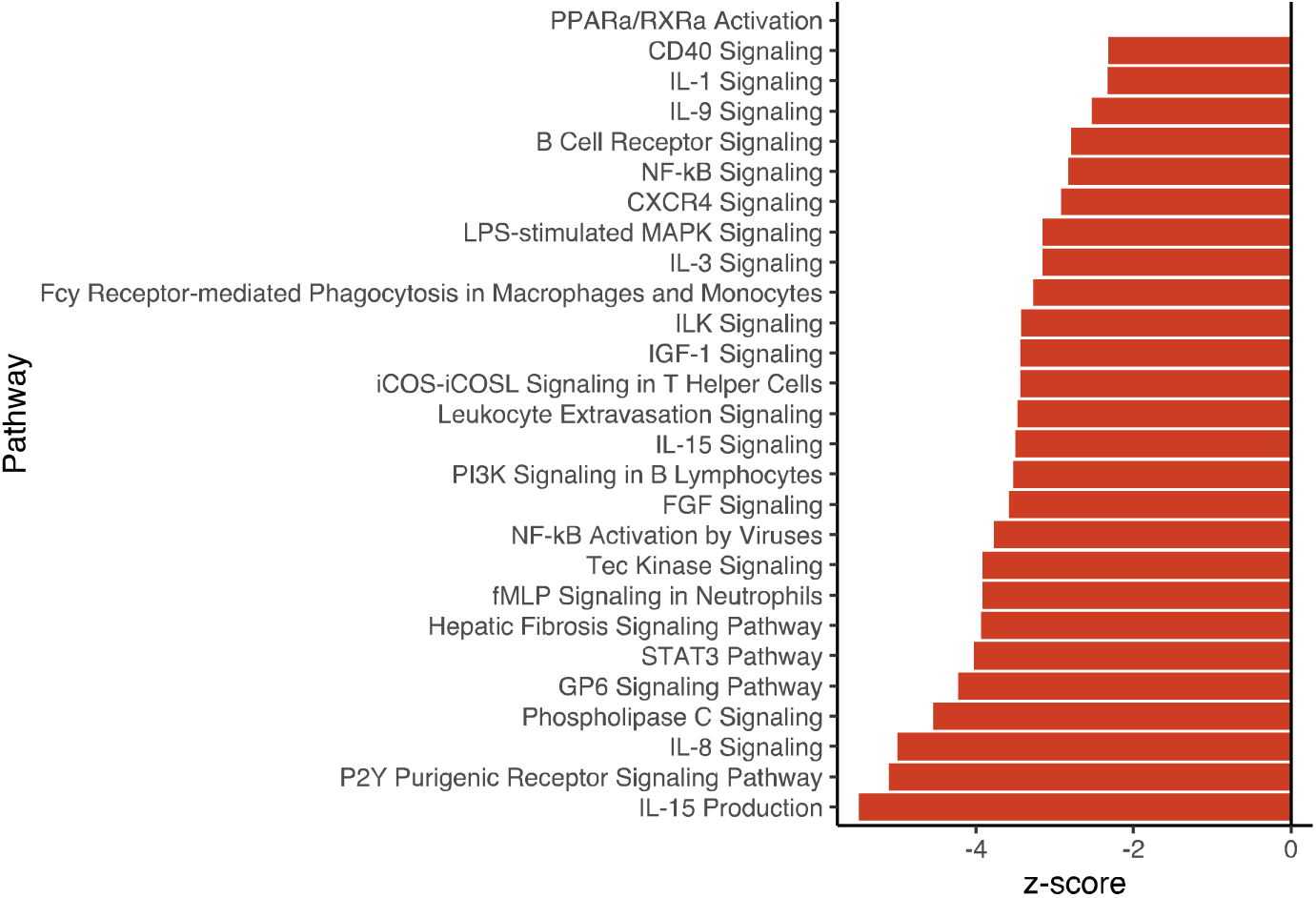
Summary of biological pathways involved in immunity that were significantly activated/inactivated as a result of infection.

Analysis of upstream regulators also demonstrated multi-dimensional responses to infection (121 regulators significant affected; 27 with roles in immunity; **Supplementary Figure 1**, **Supplementary File 3**). Similar to pathway results, regulators displayed broad patterns of immune suppression. Anti-fibrotic regulator heptatic nuclear factor 4-alpha, HNF4A (z-score = 1.152, *p_adj_* = 6.06E-7; (Yue et al., 2010) was significantly activated as a result of infection. Several important adaptive immune components such as interleukin-4 (IL4; Heeb et al., 2020) and IgE complex (Galli and Tsai, 2012) were also suppressed in infected fish. Together, gene expression and pathway analyses reveal broad immune suppression associated with infection of *G. aculeatus* with *S. solidus*. This suppression in infected fish could arise from two processes: initially immune-suppressed fish might have been more vulnerable to infection, or the successfully established parasites may be actively suppressing host immunity. Time-series analyses of individual host expression could distinguish these alternatives, but is not practical because fish must be euthanized to obtain head kidney samples.

### Comparing cestode-induced expression to previous results

Interestingly our results showed little overlap with those from previous transcriptomic study of the response to *S. solidus* infection in pure-cross families (as opposed to hybrids) of these same stickleback populations (Lohman et al., 2017). Only 14 genes were shared between our results and theirs; all but four responded in similar directions across both studies (**Table 1**). Two of these shared genes of particular interest were: interleukin 8 and fibronectin. Infection was associated with higher expression of the extracellular matrix glycoprotein, fibronectin. This is potentially indicative of induction of fibrotic resistance phenotypes, as fibronectin production is associated with increased fibrosis (Duffield et al., 2013), which can encapsulate the parasite in a web of scar tissue that limits its movement and foraging. In contrast, interleukin 8, an important immune chemokine, showed slightly disparate patterns of expression across the two studies. This transcript increased in expression in all infected fish in our study, but in the previous study, responded differently to infection across populations. Increased expression of IL8 was associated with infection in fibrosis-prone Roberts fish, while in non-fibrotic Gosling fish IL8 expression was lower in infected individuals (Lohman et al., 2017). Finally, 76 of the genes that were differentially expressed in our study, were also differentially expressed in a study of liver transcriptome response to *S. solidus* infection by stickleback in Germany (Haase et al., 2016). The majority of these genes (~68%; *p* = 0.001954, χ^2^ = 9.5921, df = 1) responded in similar directions (**Supplementary File 4**). Interestingly, these two studies used *G. aculeatus* and *S. solidus* from different continents, suggesting long-term conservation of infection response of *G. aculeatus* to *S. solidus*, and consistency across tissue types. Increasing study of host-parasite dynamics across the circum-polar range of stickleback will aid in refining a list of widely shared mechanisms of infection response in this system.

**Table 1:** Comparison of infection-associated significantly differentially expressed genes to results from a previous study (Lohman et al., 2017) using the same two source populations.

| Transcript ID | Annotation | Current Study |  | Lohman et al. |  |  |
| --- | --- | --- | --- | --- | --- | --- |
| | | LFC | $p_{adj}$ | LFC | $p_{adj}$ | Factor |
| ENSGACT00000014173.1 | Annexin A2-A | 0.811 | 0.00350 | 1.265 | 0.0730 | Infection |
| ENSGACT00000015567.1 | Chromobox protein homolog 8 | -0.367 | 0.0210 | -0.961 | 0.00699 | Infection |
| ENSGACT00000023042.1 | Dopamine beta-hydroxylase | -0.466 | 0.0698 | -2.170 | 0.0662 | Infection |
| ENSGACT00000020041.1 | Fibronectin | 1.060 | 3.01E-12 | 0.887 | 0.0662 | Infection |
| ENSGACT00000004524.1 | Glycine--tRNA ligase | 0.235 | 0.0205 | 0.592 | 0.0638 | Infection |
| ENSGACT00000013702.1 | Guanine nucleotide-binding protein-like 3-like protein | -0.337 | 0.0637 | -0.393 | 0.0929 | Infection |
| ENSGACT00000025278.1 | Interleukin-8 | 0.462 | 5.64E-4 | 1.169 | 0.0582 | Interaction |
| ENSGACT00000015612.1 | Protein cornichon homolog 1 | 0.256 | 0.0119 | 0.455 | 0.0662 | Infection |
| ENSGACT00000008095.1 | SID1 transmembrane family member 2 | 0.262 | 0.0736 | -0.557 | 0.0662 | Infection |
| ENSGACT00000018426.1 | Sodium channel protein type 2 subunit alpha | -0.299 | 0.0938 | -1.104 | 0.0862 | Infection |
| ENSGACT00000011810.1 | Sorting nexin-3 | 0.326 | 0.00612 | 0.425 | 0.0662 | Infection |
| ENSGACT00000008510.1 | Tubulin alpha chain | 0.275 | 0.0377 | 0.978 | 0.0953 | Interaction |
| ENSGACT00000026489.1 | Unannotated | -0.441 | 0.0617 | -0.823 | 0.0638 | Infection |
| ENSGACT00000008169.1 | Unannotated | 1.415 | 2.05E-4 | 3.239 | 0.0662 | Infection |

### Variation in gene expression and immune response among cross types

The Baldwin Effect suggests that plastic traits (e.g., genes responding generically to cestod infection) should also contribute to between-population divergence (either constitutive, or in infection response). To test this, we next summarize the constitutive and infection-induced gene expression variation among three different cross types of *G. aculeatus* representing F2 backcrosses and intercrosses of fish from a resistant and a susceptible lake (Roberts and Gosling, respectively). There was considerable constitutive variation in gene expression patterns among cross types. Of the 15,354 genes tested, 11,321 varied constitutively between any two given cross types: 7745, 10601, and 1202 transcripts were differentially expressed between F2 vs GBC, F2 vs RBC, and RBC vs GBC respectively (*p_adj_* < 0.10, 10% FDR; **Figure 2; Supplementary File 1**). Of course many of these genes will have no bearing on immunity, but a large portion of these differentially expressed genes have immunological functions, corresponding to diverse components of immunity (**Table 2**).

**Figure 2:**
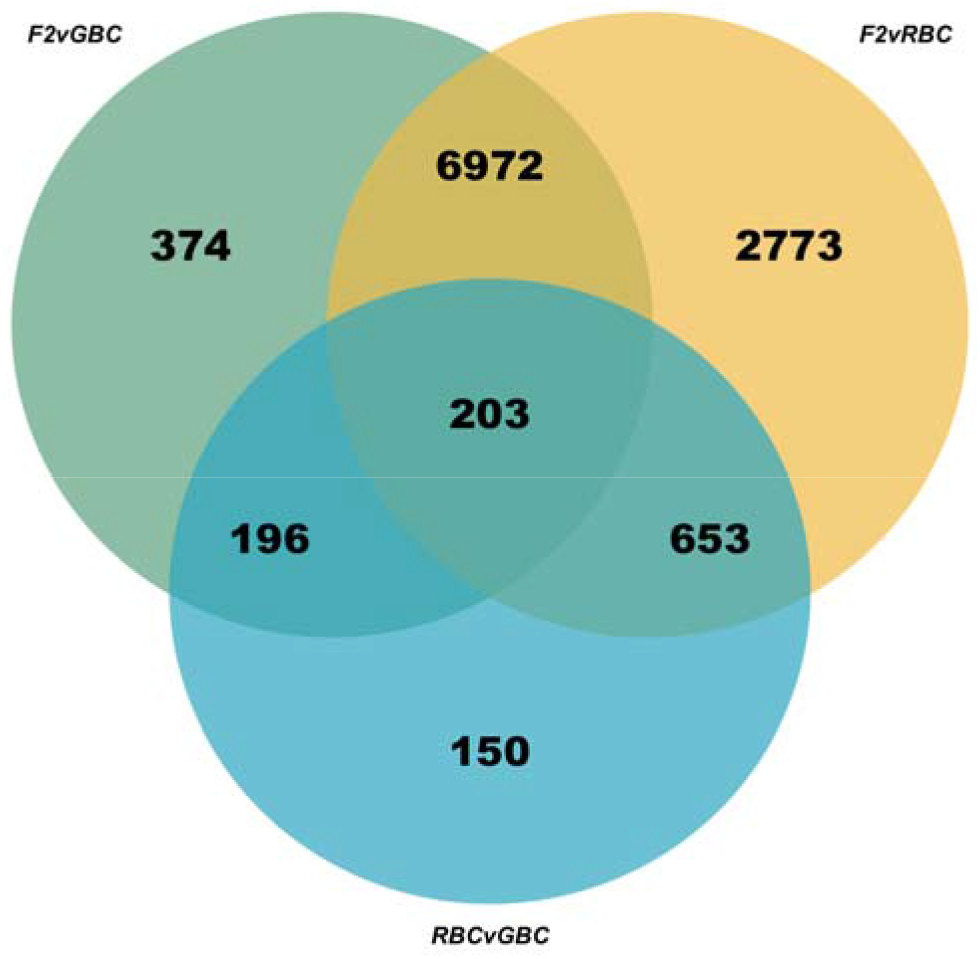
Venn diagram of overlap in significantly differentially expressed genes among all three cross type comparisons.

**Table 2:** Example list of genes differentially expressed among cross types with putative functions in immunity. A full list of differentially expressed genes for each contrast can be found in **Supplementary File 1**.

| Transcript ID | Annotation | F2 vs. GBC |  | F2 vs. RBC |  | RBC vs. GBC |  |
| --- | --- | --- | --- | --- | --- | --- | --- |
|  |  | LFC | *p<sub>adj</sub>* | LFC | *p<sub>adj</sub>* | LFC | *p<sub>adj</sub>* |
| ENSGACT00000002677.1 | B-cell receptor CD22 | 0.718 | 0.00504 | 1.189 | 1.81E-8 | -0.472 | 0.0702 |
| ENSGACT00000008607.1 | Carboxypeptidase N catalytic chain | -0.335 | 0.0551 | -0.654 | 4.93E-6 | 0.319 | 0.0848 |
| ENSGACT000000021611.1 | Complement component C8 alpha chain | 0.755 | 0.0863 | 1.558 | 1.16E-5 | -0.803 | 0.0639 |
| ENSGACT000000012995.1 | C-type lectin domain family 4 member E | 1.207 | 0.0379 | 2.348 | 5.21E-7 | -1.141 | 0.0364 |
| ENSGACT000000024032.1 | Gelsolin | -1.784 | 3.05E-9 | -0.856 | 2.16E-4 | -0.927 | 0.0974 |
| ENSGACT000000005722.1 | Granulins | 0.671 | 3.01E-4 | 0.331 | 0.0164 | 0.340 | 0.0562 |
| ENSGACT000000012867.1 | H-2 class I histocompatibility antigen, L-D alpha chain | 3.775 | 3.62E-4 | 2.026 | 0.0113 | 1.748 | 0.0798 |
| ENSGACT000000010258.1 | Histone acetyltransferase p300 | -0.477 | 0.0473 | -0.905 | 4.06E-6 | 0.428 | 0.0915 |
| ENSGACT000000001096.1 | Macrosialin | -0.506 | 0.0168 | -0.888 | 4.44E-7 | 0.382 | 0.0849 |
| ENSGACT000000002730.1 | Transforming growth factor beta activator LRRC33 | -0.616 | 0.00347 | -1.000 | 1.21E-8 | 0.383 | 0.0747 |
| ENSGACT000000024916.1 | Ubiquitin-conjugating enzyme E2 N | 0.256 | 0.0927 | -0.319 | 0.00785 | 0.575 | 5.25E-6 |

A large fraction of the infection-responsive genes (1812 of the 2369 differentially expressed genes associated with infection, or 76%) also varied in expression between two or more cross types. However the overlap between the infection-responsive and constitutively-different genes was no greater than random expectations (*p* = 0.2821, χ^2^ = 1.1569, df = 1). Furthermore, we found that effect sizes and directions of infection and genotype main effects were inversely correlated for the subset of genes significantly differentially expressed both in infected fish and between RBC vs. GBC fish (*p* = 0.0005439, df = 145, cor = −0.2818,). This would indicate that genes that respond to infection have higher baseline expression in GBC versus RBC fish, the opposite of our predictions based on the Baldwin Effect. Lower constitutive expression by RBC fish of immune responsive genes may be a result of reduced investment; resistant hosts may be successful enough at avoiding infection through fibrosis that they need not invest in other “normal” immune related genes. Alternatively, higher constitutive expression of genes which are suppressed during infection by RBC could be evidence of counter adaptation to parasite immune suppression (discussed later). Resistant fish may avoid infection by constitutively increasing expression of key immune gens that are typically targeted by parasites and subsequently suppressed. Further experimental studies will help distinguish whether either of these hypotheses is applicable to this system.

Despite the lack of evidence for the Baldwin Effect when examining the whole transcriptome, there are numerous individual genes whose expression patterns are highly consistent with the Baldwin Effect. In particular, a number of genes annotated as having key immune functions fit our expectations. For example, signal transducer and activator of transcription 3 (STAT3), is expressed higher in RBC fish than GBC fish, and significantly increased in expression as a result of infection. STAT3 is a multi-faceted immune cytokine which contributes to both the differentiation of IL-17 producing T cells and is also critical for CD8 T-cell memory (O'Shea et al., 2013). Based on both its well-described immune functions, and evidence presented here regarding variation of expression of this gene both during infection and across genotypes, STAT3 is an excellent candidate resistance gene in the *G. aculeatus* – *S. solidus* system. Further studies, including functionalization using transgenic methods should elucidate the mechanisms by which this gene contributes to infection response, and potentially *S. solidus* resistance.

In addition to broad variation among cross-types in gene expression, numerous pathways and upstream regulators, including many involved in immunity, were also significantly differentially expressed between crosses (*p_adj_* < 0.10, 10% FDR), independent of infection status. A total of 240, 313, and 94 pathways varied in activation state when comparing F2 vs. GBC, F2 vs. RBC, and RBC vs. GBC respectively (**Supplementary Figure 2; Supplementary File 2**). Additionally, 105, 164, and 22 upstream regulators were predicted to have differential activity between cross comparisons F2 vs. GBC, F2 vs. RBC, and RBC vs. GBC respectively (**Supplementary File 3**). Differentially expressed pathways represented many different immune components including IL-2 Signaling (**Table 3**). Interluekin-2 (IL-2) is a key immune cytokine with many roles including maintenance and differentiation of T cells (Boyman and Sprent, 2012). Several pathways were affected by both cross type and infection, including B Cell receptor signaling and NF-kB activation/signaling. B cells uniquely express both B cell receptors and Toll-like receptors, which signal using NF-kB, allowing for the linking of innate and adaptive arms of immunity (Rawlings et al., 2012). Integration of diverse components of immunity is likely key in response to *S. solidus*, though the dynamics of this cross-talk may vary among crosses. In sum, our results demonstrate significant constitutive variation among crosses, including variation in putative immune components.

**Table 3:** Example list of pathways that were differentially activated among cross types and have putative functions in immunity. A full list of differentially activated pathways for each contrast can be found in **Supplementary File 2**.

| Pathway | F2 vs. GBC |  | F2 vs. RBC |  | RBC vs. GBC |  |
| --- | --- | --- | --- | --- | --- | --- |
|  | z-score | *p*<sub>adj</sub> | z-score | *p*<sub>adj</sub> | z-score | *p*<sub>adj</sub> |
| B Cell Receptor Signaling | 2.945 | .00204 | 1.889 | 3.00E-6 | 0.816 | 0.0398 |
| FLT3 Signaling in Hematopoietic Progenitor Cells | 2.921 | 0.00398 | 2.214 | 3.50E-5 | -0.535 | 0.0324 |
| Hepatic Fibrosis Signaling Pathway | 5.031 | 0.00138 | 5.261 | 4.00E-6 | -1.677 | 0.0871 |
| IGF-1 Signaling | 2.121 | 0.00407 | 1.852 | 4.70E-5 | 0 | 0.0912 |
| IL-2 Signaling | 1.414 | 0.0589 | 0.816 | 0.00372 | -0.378 | 0.0871 |
| IL-6 Signaling | 1.761 | 0.0275 | 1.387 | 0.00105 | -0.728 | 0.0776 |
| Leukocyte Extravasation Signaling | 3.960 | 0.0186 | 3.761 | 4.00E-6 | 0.200 | 0.0537 |
| Lymphotoxin $\beta$ Receptor Signaling | 2.668 | 0.00708 | 1.225 | 9.80E-5 | -0.378 | 0.0537 |
| NF-κB Activation by Viruses | 2.600 | 0.0832 | 2.03 | 3.10E-5 | -0.277 | 0.0479 |
| NF-κB Signaling | 3.501 | 0.0427 | 3.064 | 3.39E-4 | -1.091 | 0.0977 |
| P2Y Purigenic Receptor Signaling Pathway | 3.317 | 1.62E-4 | 2.994 | 2.24E-7 | -1.213 | 0.0324 |

There is also appreciable overlap between the infection-induced pathways, and the pathways that diverge constitutively between populations. Of the 169 pathways significantly differentially activated as a result of infection, 158 were also differentially activated among two or more cross types (*p* < 0.001, χ^2^ = 21.528, df = 1). Thus, while our results do not support the Baldwin Effect on the level of individual genes (as described in the preceding section), there is evidence supporting this theory at the pathway level. Further investigation of host immune response to parasite at multiple levels (gene, regulatory network, whole pathway) will improve understanding of the evolutionary patterns driving variation at each level.

### Interactive effects of cross-type and infection response

Previous analysis of pure F1 ROB versus GOS fish showed that transcripts with main-effects of infection (shared across all genotypes) vastly outnumbered transcripts with genotype-specific responses to infection (genotype*infection interactions; (Lohman et al., 2017). Here we report similar results. A total of 569 genes exhibited interactions between cross type and infection, however the vast majority of these were only significant when considering differences in response to infection between F2 fish and GBC fish (559/569). Sixty-four of these genes which responded differentially to infection in F2 vs. GBC fish have potential roles in immunity or fibrosis, including 5 collagen transcripts (**Supplementary File 1**). In contrast, few genes responded differentially to infection when comparing F2 vs. RBC and RBC vs. GBC fish. Thirteen transcripts responded differentially to infection in F2 vs. RBC fish, most of which did not have function in immunity/fibrosis. Finally, examination of differences in response to infection between the two most disparate crosses (RBC/GBC) provides some insight regarding potential candidate genes contributing to resistance (**Figure 3**). Four genes responded differentially to infection between these crosses, two of which had roles in immunity: CCN family member 3 (CCN3) and SH2 domain-containing protein 1A (SH2D1A). CCN3 expression suppresses fibrosis responses through the modification of expression of other CCN family proteins (Abd El Kader et al., 2013). CCN3 decreased more significantly in response to infection in resistant (RBC) fish, perhaps contributing to observed fibrosis phenotypes. SH2D1A is an important mediator of humoral immunity, particularly long-term immune memory, largely through regulation of CD4^+^ T cell functioning (Crotty et al., 2003). Furthermore, SH2D1A may affect NKT cell ontogeny (Nichols et al., 2005) and differentiation of T_H_2 cells (Wu et al., 2001). Interestingly, expression of this key component of adaptive immunity was maintained in resistant fish (RBC) in response to infection, but significantly decreased in the susceptible cross (GBC), highlighting the importance of adaptive immunity, and potentially long-term immune memory, in resistance to cestode infection in *G. aculeatus*.

**Figure 3:**
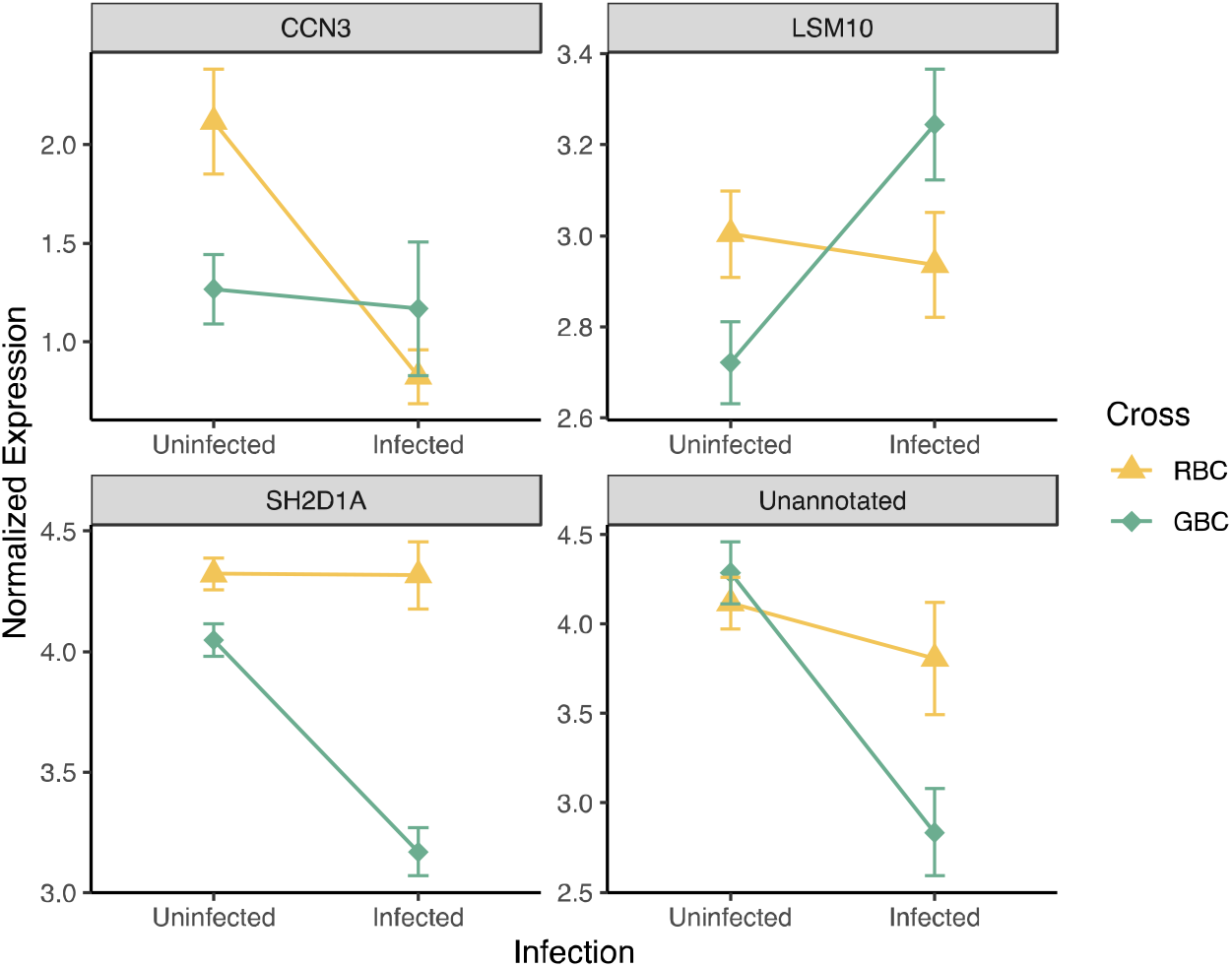
Interaction plot displaying changes in expression of the four genes that responded differentially to infection in RBC vs. GBC fish.

In sum, our results indicate that while crosses respond roughly equivalently to infection, there are many constitutive differences among these crosses, including in expression of immune genes. These differences may underlie population differences in parasite resistance. However, while there is overlap between the induced and constitutive effects, at a whole-transcriptome gene-level scale this overlap is no more extensive than null expectations. Thus our study does not provide significant support for the applicability of the Baldwin Effect to the evolution of host resistance in this system, at the whole-transcriptome, individual gene level. Furthermore, the Baldwin Effect postulates that beneficial traits (i.e. parasite resistance) may first arise as plastic responses, which are selected for, eventually resulting in canalization of these inducible differences. Our results also do not show strong evidence for either variation in plastic responses to parasites, as we observed few infection by genotype interactions. Instead we see a mix of these two patterns: many genes that are responsive to parasites do vary among cross-types, but not a statistically significant excess proportion. Furthermore, some genes do show variation in plasticity (i.e response to infection among crosses), however such interaction effects occur for only a small proportion of genes (four total). However, our results are inconclusive regarding whether this plasticity is stronger in GBC fish compared to RBC fish across these 4 genes, likely due to the small sample size (one-tailed Fisher’s exact test *p* = 0.07143). Thus our findings suggest that evolution of host resistance in the *G. aculeatus* – *S. solidus* is likely the result of a combination of variation in both constitutive and inducible responses.

### Expression changes associated with fibrosis

In addition to evaluating the relevance of the Baldwin Effect to evolution of host resistance, our findings also shed light on a putative resistance phenotype in this system: fibrosis. Fibrosis is a common immune pathology across vertebrates (Wick et al., 2010;Sgalla et al., 2016;Vrtílek and Bolnick, 2020), frequently associated with parasitic infections (Wynn et al., 2004;Wilson et al., 2007;Niu et al., 2019). Often excessive fibrotic responses can cause health issues, including in humans (Friedman, 2004;Wynn, 2008;Todd et al., 2012). In *G. aculeatus*, recent study has demonstrated that fibrosis is an induced response to *S. solidus* in some stickleback populations, and is associated with reduced cestode growth (Hund et al., 2020; Weber et al. *in prep*). More broadly, teleost fish in general are almost all susceptible to peritoneal fibrosis in response to an immune challenge (Vrtílek and Bolnick, 2020). Our data identified strong transcriptomic signatures of fibrosis: 5,825 genes were differentially expressed in fibrotic fish compared to those not displaying the fibrosis phenotype. Many of these genes have putative roles in immunity and fibrosis, including 22 genes involved in complement activation. Numerous previous studies have indicated potential cross-talk between components of the complement cascade and fibrosis responses (Xavier et al., 2017;Liu et al., 2018). Similar to the effects of infection, the majority of these genes were downregulated in fibrotic fish. We also observed 36 collagen genes that were differentially expressed in fibrotic fish, almost all of which were also downregulated. Fibrotic tissue is formed via the deposition of collagen and other extracellular matrix components (Wynn, 2008). The large-scale down-regulation of these complement and collagen genes, as well as other immune and fibrosis-related transcripts, in fibrotic fish may be an artifact of the time of sampling: 42 days after exposure. It is possible that these genes are activated earlier on in the infection time course, but have since been downregulated.

Pathway and upstream regulator analyses also revealed significant down-regulation of immune-related processes in fibrotic fish (**Figure 4; Supplementary Figure 1**). Many (257) biological pathways were significantly differentially activated in fibrotic fish, 53 of which are related to immunity, defense, or fibrosis responses. Most of these immune-related pathways were also significantly differentially activated in fish infected with *S. solidus*. However these shared pathways were almost always downregulated to a greater degree in infected fish, compared to fibrotic fish (**Figure 5**). Many of these shared pathways have potential ties to fibrotic processes, including hepatic fibrosis signaling pathway and IGF-1 signaling. IGF-1 stimulates the differentiation of fibroblasts to promote a potent fibrosis response (Hung et al., 2013). Other pathways uniquely activated in fibrotic fish include the key innate immune component, Toll-like receptor signaling, which was activated in fibrotic fish. Numerous TLR receptors can bind to parasite antigens to activate an immune response (Campos et al., 2001;Coban et al., 2005), though their links to fibrotic responses are unclear. Finally, pathways related to NFAT regulation of immune response were significantly downregulated in fibrotic fish. NFAT contributes to the regulation of T cell development, diversification, and activation and affect T-cell lineage differentiation (i.e. T_H_1 vs. T_H_2; Macian, 2005). Down regulation of this pathway suggests fibrotic fish may have reduced regulation of T-cell activity, allowing for more potent responses to *S. solidus*. In sum, pathway analyses suggest that, compared to infected fish without fibrosis, fibrotic fish have reduced down regulation of immune pathways, and activation of unique pro-defense pathways, which may allow for the induction of these putative resistance responses. Upstream regulators show similar patterns. While several key immune regulators were downregulated in both infected and fibrotic fish, fibrotic fish also demonstrated unique signatures of immune response. For example, negative regulator of immunity such as HDAC4 (Yang et al., 2019) was uniquely suppressed in fibrotic fish. Furthermore, anti-fibrotic regulator heptatic nuclear factor 4-alpha, HNF4A (Yue et al., 2010) was significantly activated as a result of infection, but suppressed in infected fish. Combined these results suggest that while infection is marked by a general reduction in immune pathways, this suppression is weaker or absent in fibrotic fish that more effectively reduce cestode growth.

**Figure 4:**
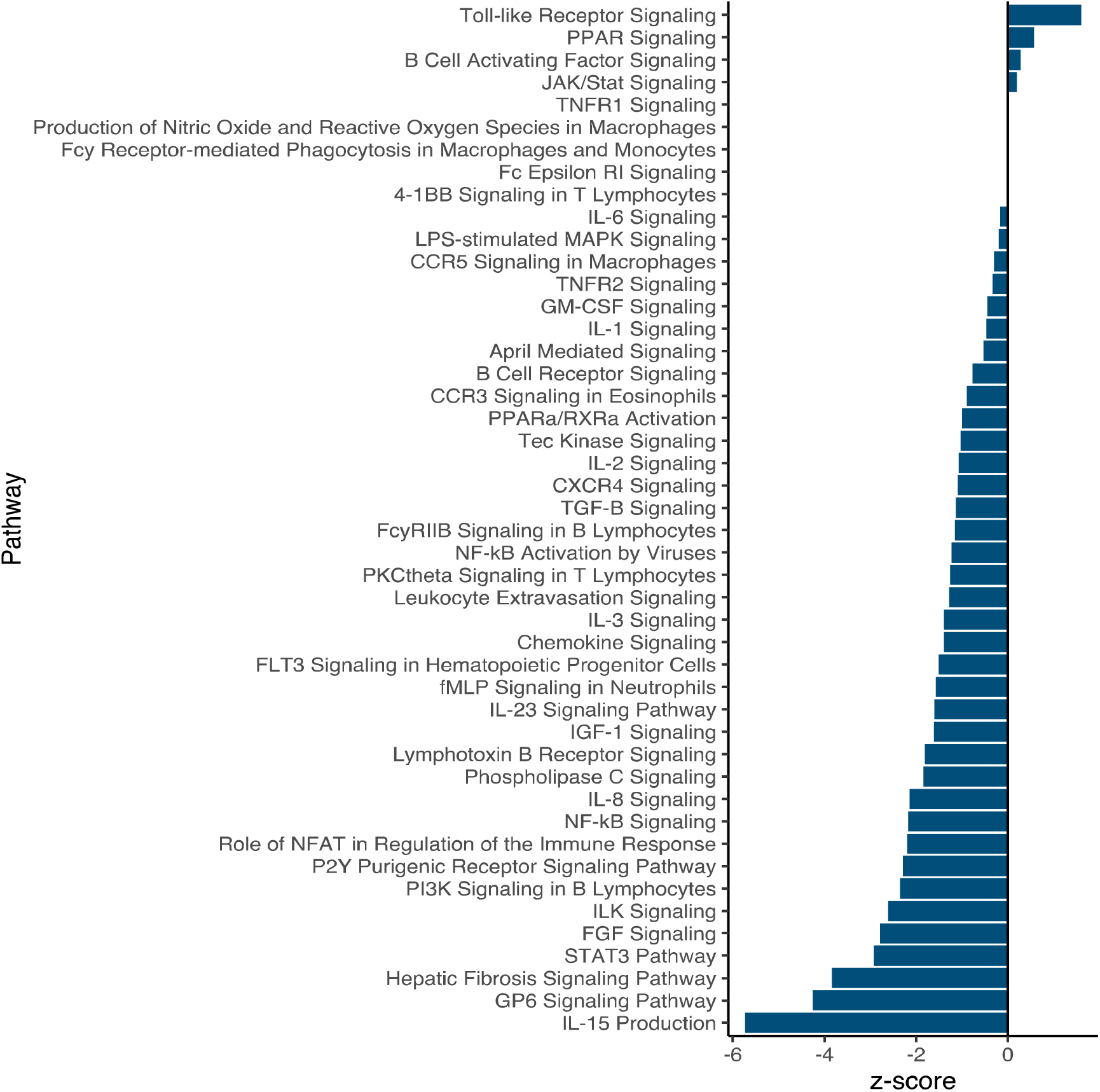
Summary of biological pathways involved in immunity that were significantly activated/inactivated as a result of fibrosis.

**Figure 5:**
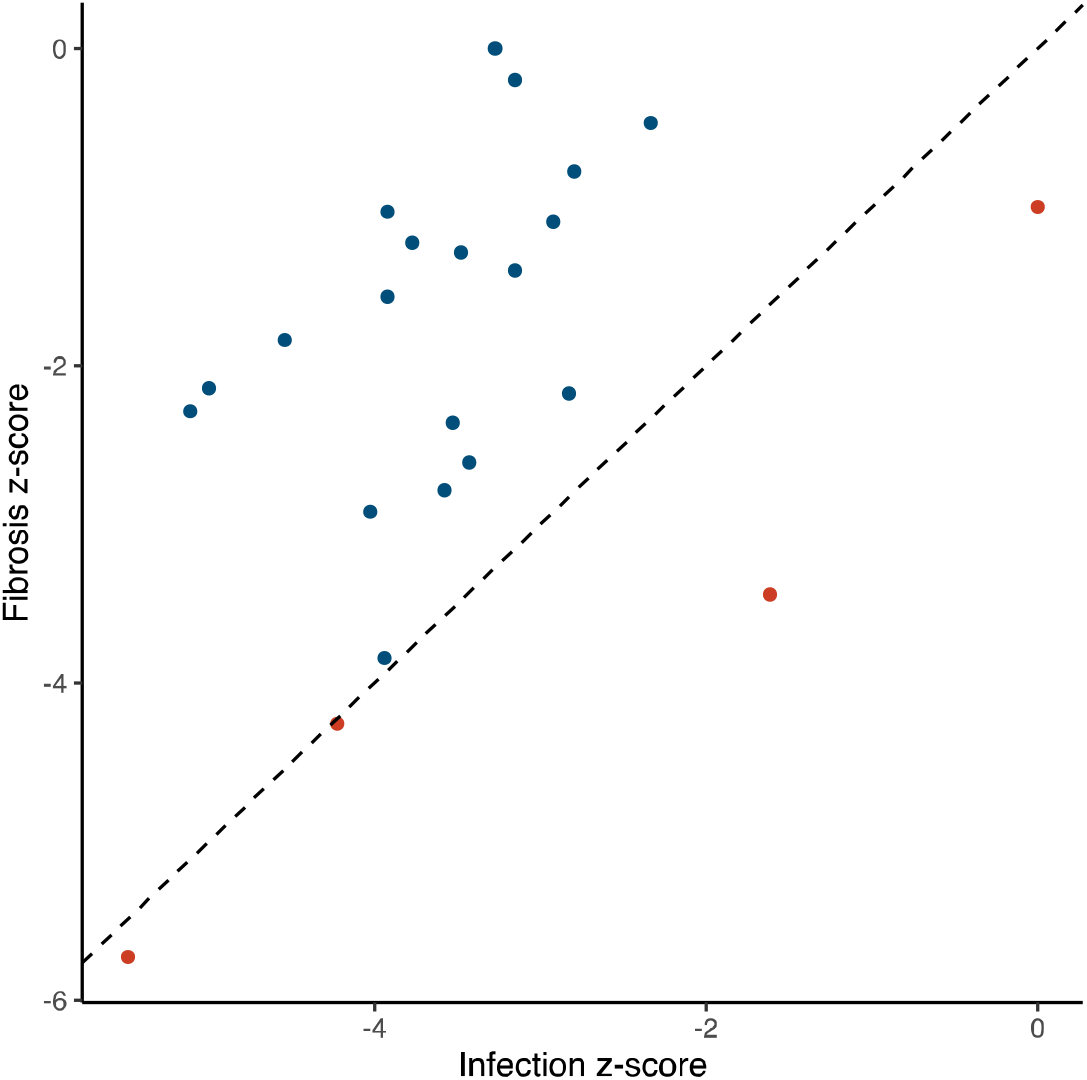
Comparison of activation (z-score) of immune-related pathways that were significantly activated/inactivated as a result of both fibrosis and infection. Dotted line represents equivalent activation as a result of both factors; points in blue are more activated in fibrotic fish; points in red are more activated in fibrotic fish.

The association between infection and immunosuppression is particularly interesting as immunosuppression is widely known in helminths in general (Maizels et al., 2004;Maizels and McSorley, 2016;Maizels et al., 2018), and has been documented for *S. solidus* in particular (Scharsack et al., 2004). Thus it is possible the described patterns are indicative of cestode suppression of host immune response during infection. In contrast, fibrotic fish show weaker signatures of immune reduction, and in some instances demonstrate patterns of expression opposite to those displayed by infected, non-fibrotic fish. Furthermore, these specific pathways and genes that demonstrate disparate patterns between fibrotic and infected-but-nonfibrotic fish (e.x. HNF4A), are also highly variable among cross-types. Combined, these analyses suggest that cestodes may act to suppress immune responses in their host, but that this immunosuppression differs between host genotypes, and between fibrotic and non-fibrotic fish. These findings suggest that evolution of resistance may also be dependent on the acquisition of traits to overcome or avoid parasite manipulation of host immunity.

## CONCLUSIONS

Despite broad interest in host-parasite interactions and associated immune responses, relatively few studies have identified the mechanistic immunogenetic basis of rapid microevolution of vertebrate host immunity in the wild. Current knowledge of these processes are limited to a few natural examples, such as the rapid evolution of polygenic resistance to myxomatosis in rabbits, and selection for resistance to nematode parasites in Soay sheep (Hayward et al., 2011). Here we contribute a new, non-mammalian example of natural evolution of parasite resistance. We leveraged a large transcriptomic data set to evaluate the relevance of one evolutionary theory, the Baldwin Effect, to the recent evolution of host resistance. First, we characterized the general response to a parasite in the well-studied *G. aculeatus* – *S. solidus* host-parasite system. Our results showed that fish mount a diverse immune response to parasitic infection, involving multiple genes and pathways. Then, we compared both infection-induced responses and constitutive differences among fish cross types with variable parasite resistance. Interestingly, cross types showed mostly similar induced responses to parasites (e.g., little evidence for canalization, the loss of expression plasticity in one population). Instead, we demonstrate patterns of constitutive variation among crosses. Although these constitutive differences overlap with many genes that change plastically with infection, that overlap is not significantly greater than null expectations. Thus our results fail to provide conclusive support for the applicability of the Baldwin Effect to evolution of host resistance to *S. solidus*, at least at the whole-transcriptome scale. Further investigation in diverse systems, as well as experimentation with rapidly reproducing host-parasite models will be useful in clarifying the roles of constitutive vs. inducible variation in evolution of host resistance to parasites. Also, the Baldwin Effect may be appropriate to describe specific immune genes or pathways, even if not the entire transcriptomic response to infection. Additionally, our results identify transcriptomic signatures contributing to a putative resistance phenotype, fibrosis. Not only do we identify key genes and pathways associated with this response, but we also provide evidence that fibrotic (i.e. resistant) fish apparently are refractory to immune suppression associated with infection in other fish. This suggests host evolution of counter mechanisms may also be key in the evolution of resistance. Combined, our results highlight new potential theories regarding the patterns driving evolution of resistance in host-parasite systems.

**Table 4:** List of genes that were significantly differentially expressed as a result of the interaction between cross type (RBC vs. GBC) and infection.

| Transcript ID | Annotation | RBC vs. GBC |  |
| --- | --- | --- | --- |
| | | LFC | $p_{adj}$ |
| ENSGACT00000015170.1 | CCN family member 3 | 4.866 | 1.45E-6 |
| ENSGACT00000018024.1 | Unannotated | -2.159 | 0.0986 |
| ENSGACT00000024555.1 | SH2 domain-containing protein 1A | -1.202 | 0.0473 |
| ENSGACT00000003040.1 | U7 snRNA-associated Sm-like protein LSm10 | 1.157 | 0.0473 |

## Supplementary Materials

**Supplementary File 1**: List of transcripts which were significantly differentially expressed in infected fish, or between 2 of the host cross types. Data includes annotation, log2fold change value, and adjusted p-value for each transcript.

**Supplementary File 2**: List of pathways which were predicted to be significantly differentially activated in infected fish, or between 2 of the host cross types. Data includes adjusted p-value, z-score (metric of activation), and included molecules for each pathway.

**Supplementary File 3**: List of upstream regulators which were predicted to be significantly differentially activated in infected fish, or between 2 of the host cross types. Data includes expression, adjusted p-value, molecule type, predicted activation, z-score (metric of activation), and affected molecules for each regulator.

**Supplementary File 4**: Genes that were commonly differentially expressed in infected fish across our study and results from a previous study of *G. aculeatus* and *S. solidus* (Haase et al. 2016).

**Supplementary Figure 1:** Bar graph displaying predicted patterns of activation for upstream regulators which were significantly differentially activated in **A)** infected, **B)** fibrotic, and **C)** both infected and fibrotic fish.

**Supplementary Figure 2**: Venn diagram of overlap in pathways which were differentially activated between each of the three cross types.

