## Supplementary figures and images for "A test of the Baldwin Effect: Differences in both constitutive expression and inducible responses to parasites underlie variation in host response to a parasite"

### Supplementary Figure 1

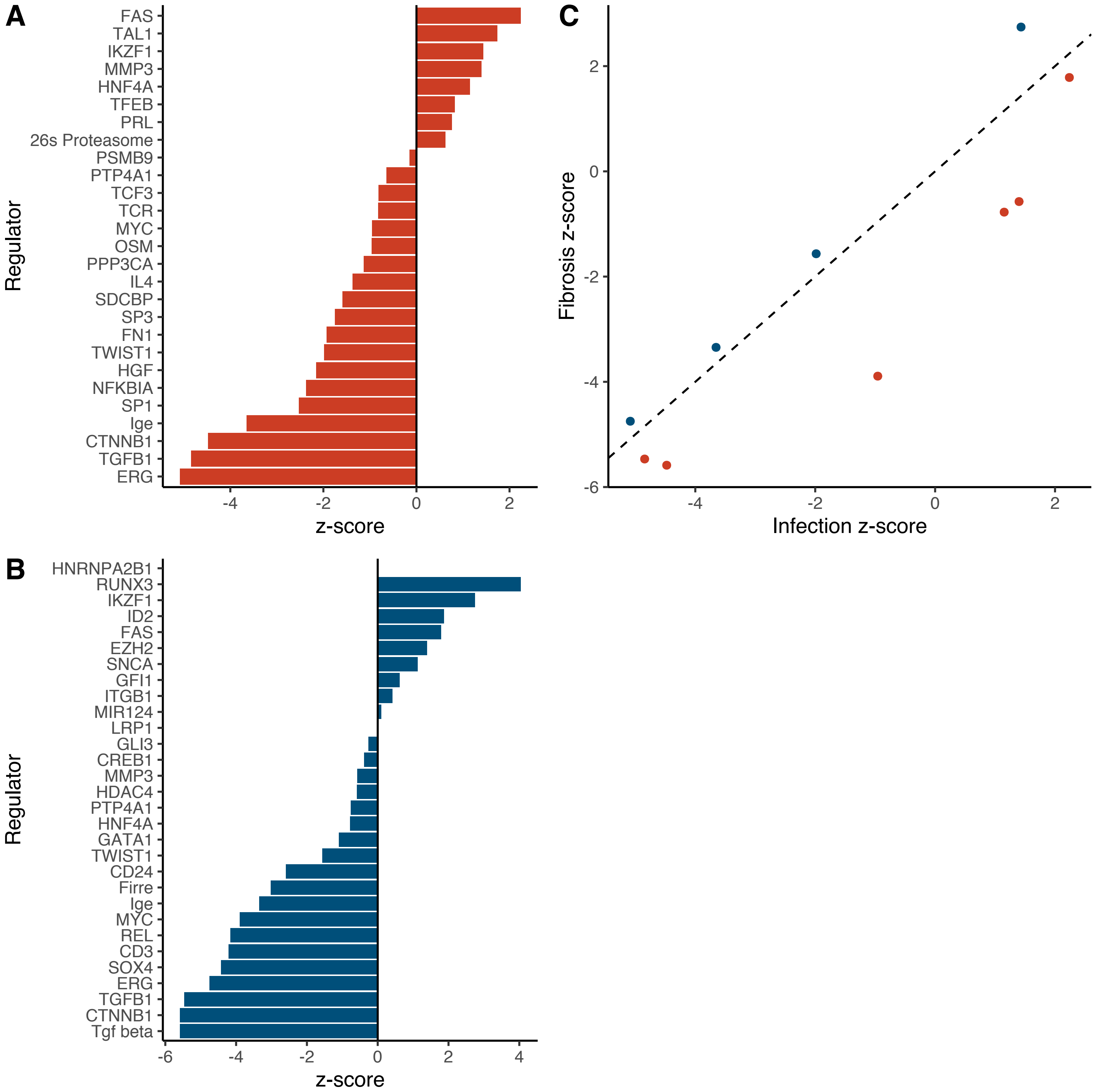

### Supplementary Figure 2

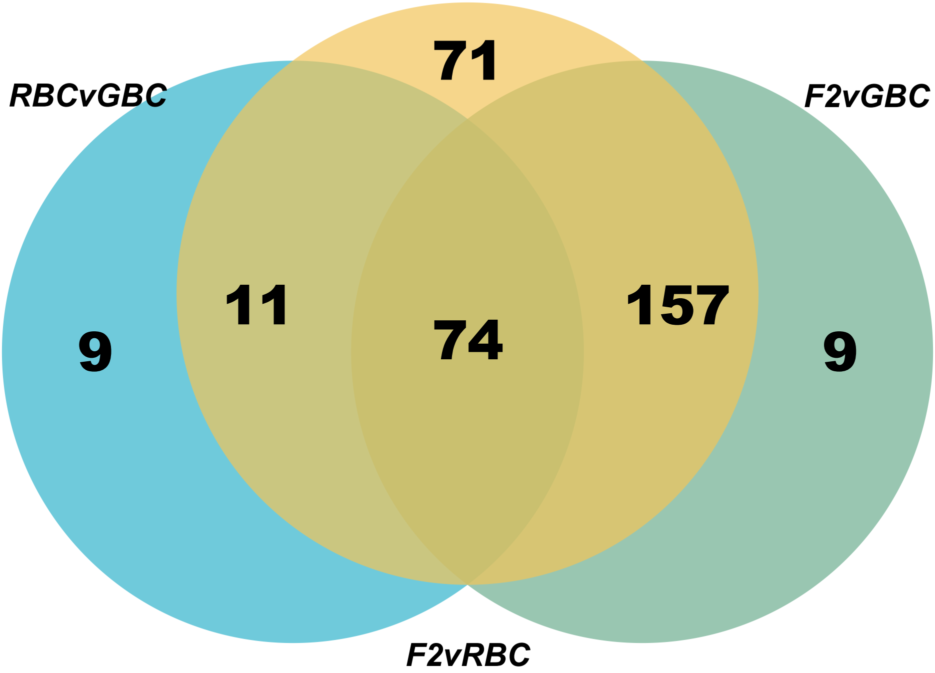
